# A multi-component power-law penalty corrects distance bias in single-cell co-accessibility and deep-learning chromatin interaction predictions

**DOI:** 10.1101/2025.08.20.671329

**Authors:** Luca Schlegel, Fabio Gómez Cano, Alexandre P. Marand, Frank Johannes

**Affiliations:** Plant Epigenomics, TUM School of Life Sciences Weihenstephan, Technical University of Munich, Munich, Germany; Department of Molecular, Cellular, and Developmental Biology, University of Michigan, Ann Arbor, USA

## Abstract

Scalable proxies for 3D genome contacts - such as single-cell co-accessibility and deep learning predictions - have emerged as powerful alternatives to chromatin capture-based methods, but predictions systematically overestimate long-range interactions. Here we show how to correct this bias using distance-based penalty functions informed by Gaussian mixture modeling and polymer-physics scaling. Using Hi-C datasets from maize, rice, and soybean, we derive tissue-specific and global consensus penalties parameterized by multi-regime power-law exponents. Applying these corrections to scATAC-seq co-accessibility scores improves their distance profiles in concordance with Hi-C and reduces long-range false positives by an average of 73% with tissue-specific penalties and 66% with the global consensus. We provide open-source code and fitted parameters to support adoption in maize, rice, and soybean.

## 1 Introduction

The three-dimensional (3D) organization of chromatin is central to gene regulation and genome stability in eukaryotes [1, 2, 3]. At fine scales, chromatin loops bring distal regulatory elements, such as enhancers and promoters, into spatial proximity [4, 5, 6]. In many plant species, these structures are embedded within larger features, including topologically associating domains (TADs) [7, 8, 9] and active/inactive (A/B) compartments [10, 1, 11, 12, 13]. While Hi-C has been instrumental in mapping this hierarchy [10], its routine application across many genotypes and tissues remains limited by cost and logistics [14].

Scalable proxy methods, including co-accessibility scores from single-cell ATAC-seq [15] or from deep learning [16], have recently emerged as powerful alternatives. Co-accessibility inferred from single-cell ATAC-seq (scATAC-seq) estimates contacts by identifying pairs of loci whose accessibility covaries across cells [17, 18]. Similarly, deep learning (DL) models learn sequence and epigenomic correlates of interaction frequency from experimental training data [19, 20]. Both approaches enable high-throughput generation of interaction-like scores across diverse tissues and species.

A key limitation is that both approaches tend to inflate long-range contacts [21, 22, 23, 24]. For co-accessibility, correlations can arise from shared regulation even when loci are not physically proximate [25]. For DL models, predictive signals can be learned without explicitly enforcing the expected distance dependence [19, 21, 26]. This behavior conflicts with the empirically observed distance-decay of contact probability and contributes to very high false positive rates at large separations; in raw crop scATAC-seq co-accessibility, long-range FPR can exceed 0.90 (Supplementary Table S2). While heuristic normalizations exist [27], a broadly usable, data-driven correction that restores distance behavior to Hi-C-like profiles has been lacking.

Here we develop a correction framework that learns species-specific distance penalties from Hi-C and applies them to proxy interaction scores (co-accessibility and DL outputs). Empirical Hi-C distance-decay is often multi-regime (piecewise scaling), so a single monotonic penalty is insufficient (see Supplementary Fig. S1). We therefore estimate multi-regime power-law exponents using Gaussian mixture modeling and use these to define tissue-specific and global consensus penalties - derived from an equally weighted aggregation of Hi-C datasets across multiple tissues and conditions - that can be applied to new datasets for each species (see Figure 1A).

**Figure 1.**
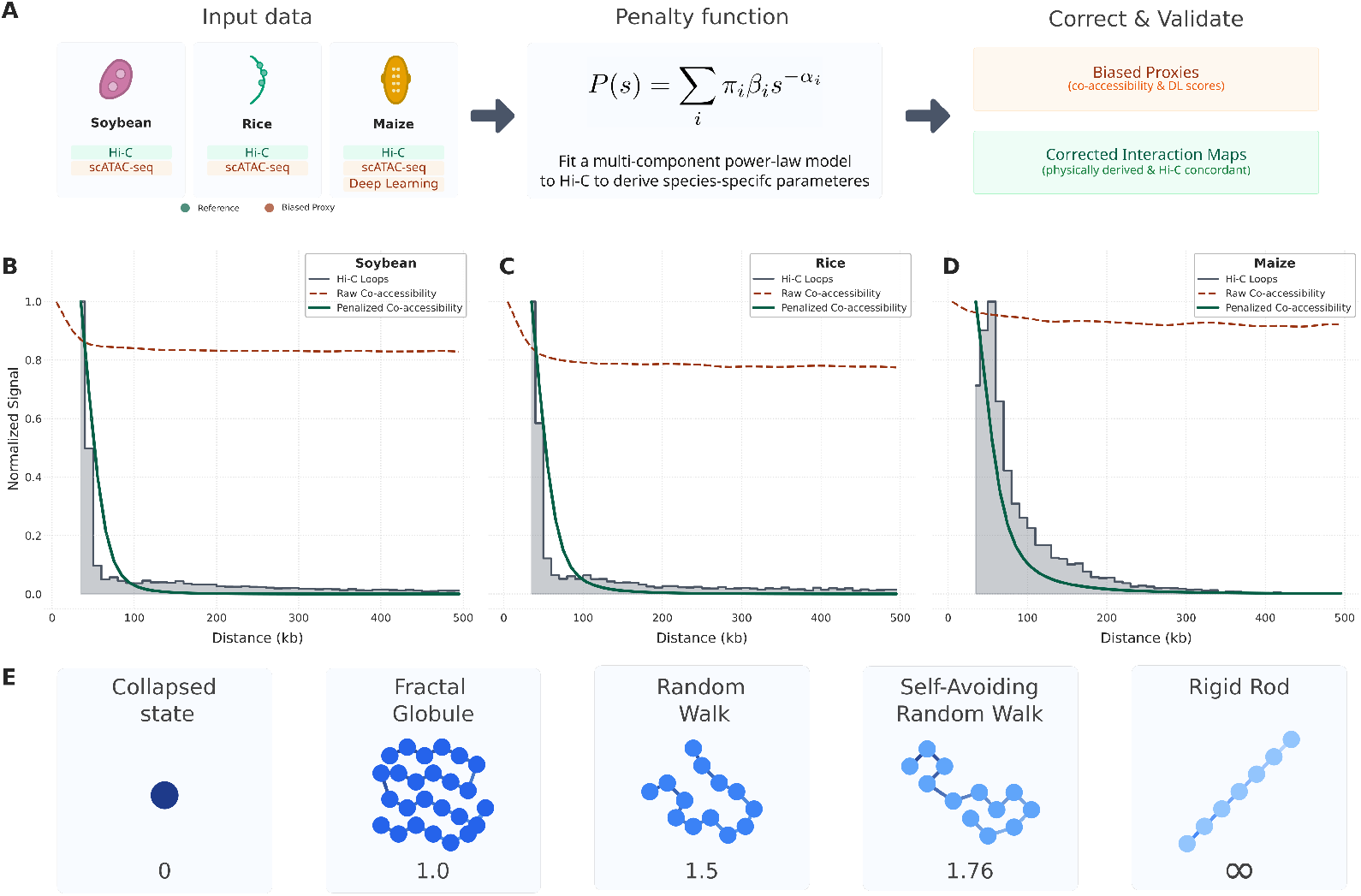
A data-driven penalty corrects distance bias in co-accessibility scores. **A)** Conceptual workflow: Multi-component power-law models fitted to species-specific Hi-C data derive penalty functions that correct biased proxies by down-weighting long-range interactions. **B–D)** Chromatin interaction profiles for soybean, rice, and maize. Hi-C decay (grey histogram) shows the expected decrease in interaction frequency with genomic distance, while raw co-accessibility scores between Accessible Chromatin Regions (ACRs) remain largely distance-independent (dark red dashed line). The GMM-based penalty corrects this bias, producing penalized profiles (dark green solid line) that closely match the Hi-C reference. All profiles are normalized to 1.0; Hi-C loop data is restricted to interactions *>* 35 kb. **E)** The power-law decay exponent *α* corresponds to distinct theoretical polymer configurations. These idealized models provide a framework for interpreting the varying decay exponents *α*_*i*_ and their regimes.

## 2 Methods

### 2.1 Data sources

All Hi-C and single-cell ATAC sequencing (scATAC-seq) datasets used in this study were publicly available and re-analyzed for consistent comparison. We obtained data from multiple tissues, developmental stages, and stress conditions. For soybean (*Glycine max*), we considered leaf tissue as well as seedling cotyledons, hooks, and hypocotyls under varying light conditions [18, 28]. For rice (*Oryza sativa*), we utilized data from 10-day-old seedlings, vegetative mesophyll, and reproductive endosperm [29, 30]. For maize (*Zea mays*), we considered leaf tissue from 6-day-old seedlings alongside reproductive tissues such as tassels and ears [27, 31]. Raw Hi-C data were sourced from GEO, SRA, NCBI BioProject (PRJNA), and GSA databases. A full list of accession numbers, key tissue information, and original publication references is provided in Supplementary Table S3. Accessible chromatin region (ACR) is used throughout to denote a peak-defined accessible region derived from scATAC-seq data.

#### Hi-C data processing, loop calling, and consensus construction

Raw Hi-C data [28, 30, 31, 32, 13] were processed using a uniform `HiC-Pro (v3.1.0)` [33] pipeline for raw read alignment, normalization, and matrix generation with non-default settings. Specifically, reads were aligned with `Bowtie2` in two passes—global followed by local—using `--very-sensitive --score-min L,-0.6,-0.2,` with seed length -L 30 (global) and -L 20 (local). Alignments with `MAPQ < 5` were discarded. A *Dpn*II digestion model was used (ligation motif GATCGATC) with the corresponding fragment BED; *cis* contacts were defined using `MIN_CIS_DIST = 20` kb. Interaction classes were reported, and singletons, multi-mappers, and PCR duplicates were removed (`GET_ALL_INTERACTION_CLASSES = 1; RM_SINGLETON = 1; RM_MULTI = 1; RM_DUP = 1`). Valid pairs were aggregated into raw contact matrices at 5 kb, 10 kb, 20 kb, 50 kb, 100 kb, 200 kb, 500 kb, and 1 Mb (upper-triangle format). The 5 kb resolution matrices were used as the basis for all downstream distance-decay profile generation. Matrices were then ICE-normalized with up to 100 iterations, filtering low-count bins at 2% and disabling high-count filtering (`FILTER_LOW_COUNT_PERC = 0.02; FILTER_HIGH_COUNT_PERC = 0; EPS = 0.1`). Default HiC-Pro settings were used otherwise. `ValidPairs` alignments were converted to the Juicer .hic file format using a combination of `HiC-Pro` and `juicer_tools` utilities. Chromatin loops were identified from. `hic`-formatted files using `HICCUPs` from the `Juicer` suite [34] of Hi-C tools with non-default parameters (`-m 512 -r 1000,2000,5000,1000 -f 0.5,0.5,0.5,0.5-p 10,8,4,2 -i 10,8,7,5 -d 5000,10000,20000,20000 --ignore_sparsity`). Loop coordinates (BEDPE) were used for distance-decay modeling. For species-level global consensus datasets, unique loop sets were pooled across tissues and developmental stages, with each dataset equally weighted in the final aggregate interaction pool.

#### scATAC-seq data processing and co-accessibility

We re-analyzed scATAC-seq data from corresponding leaf tissues for soybean [18], rice [29], nd maize [27]. ACRs were identified from pseudo-bulk signals and peak calling; a cell-by-peak binary accessibility matrix was then constructed. Co-accessibility scores between ACR pairs (separated by 2 kb to 500 kb) were calculated as previously described [21], using pseudocells generated via a *k*-nearest neighbors strategy [27] to mitigate single-cell sparsity. We retained only positively co-accessible pairs for downstream analysis. Scores represent Spearman correlation coefficients adjusted for technical confounders such as read depth. Genic and proximal ACRs were defined as regions overlapping (*>* 1 bp) gene bodies and located within 2 kb of transcription start sites (TSS) and 500 bp of transcription termination sites (TTS), respectively. Remaining ACRs were classified as distal. For co-expression analyses, pairwise Pearson correlations were calculated among genes expressed (TPM *>* 1) in at least two independent samples from the maize expression atlas [35].

### 2.2 Derivation of the distance-based penalty function

#### Data transformation for segmented power-law analysis

Chromatin loop distances were filtered to a maximum genomic separation of 5 Mb to ensure parameter stability and consistent binning. To characterize the heterogeneous nature of contact decay, loop counts were logarithmically binned (150 bins per dataset) using geometric bin midpoints to maintain accuracy in log-space transformations. Both the binned counts and their corresponding distances were transformed into log_10_ space, where the power-law relationship *P* (*s*) ∝ *s*^−*α*^ is represented as a linear function with a slope of −*α*:

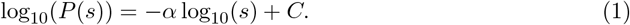

This scaling relationship, particularly for the collapsed and dense states characteristic of chromatin organization, served as the theoretical inspiration for our data-driven fitting method [36, 37].

#### GMM-based decomposition of interaction profiles

Because empirical contact profiles were frequently non-linear across the full 5 Mb range, we employed a segmented mixture-of-regressions strategy based on Gaussian Mixture Modeling (GMM) to decompose profiles into *n* distinct scaling regimes. Specifically, we used the `sklearn.mixture.GaussianMixture` class [38] with `covariance_type=‘full’` to cluster the two-dimensional distribution of log-transformed data points. This served as a data-driven method to partition the contact points into subpopulations, each corresponding to a segment with a unique decay profile.

The optimal number of components (*n*) was determined for each dataset using a dual selection process. For each *n* ∈ [1, 10], higher complexity was accepted only if it provided a significant statistical improvement (∆BIC *>* 20, strong evidence [39]) and simultaneously introduced a new regime with a characteristic decay exponent *α* differing by at least 0.25 from previously identified components (see Supplementary Fig. S2).

#### Bootstrapping of GMM transition points

To ensure the stability and relevance of the GMM results, we bootstrapped the loop-distance distributions for species-specific global consensus datasets across 10,000 replicates. Each replicate was re-binned and re-fitted using the selected *n*-component model to derive robust parameter estimates. We reported medians and 95% confidence intervals (2.5th and 97.5th percentiles) for the transition points *T*_*i*_ and scaling exponents *α*_*i*_. These parameter distributions, which confirm the structural reliability of the identified regimes, are provided in Supplementary Table S1 and Supplementary Fig. S3.

#### Construction of the GMM density function

For each of the *n* clusters identified by the GMM, a linear regression was performed on its constituent log-transformed data points. The slope of this local linear regression, slope_*i*_, was derived directly from the GMM’s cluster parameters using the relationship

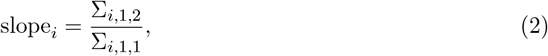

where Σ_*i*_ is the covariance matrix of the *i*-th component and Σ_*i,j,k*_ denotes the element in the *j*-th row and *k*-th column of that matrix. This slope determined the unique power-law exponent *α*_*i*_ for that segment (*α*_*i*_ = −slope_*i*_). The GMM density function *P* (*s*) is defined as

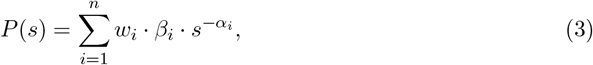

where Σ*w*_*i*_ = 1 are the weights (derived from the GMM), *β*_*i*_ are the scaling factors, and *α*_*i*_ are the fitted characteristic decay exponents. The specific sets of parameters {*α*_*i*_, *w*_*i*_, *β*_*i*_} for each species are provided in Supplementary Table S1.

### 2.3 Application and benchmarking

To correct scATAC-seq and DL scores, we derived a continuous piecewise power-law function (linear in log_10_–log_10_ space), *P*_applied_(*s*), from the GMM density function (3). Because the Hi-C loop data used for fitting was restricted to distances ≥ 35 kb, we implemented a distance plateau below this threshold where no correction is applied (*P* = 1). For all pairs with distance *s* ≥ 35 kb, the penalty is calculated as

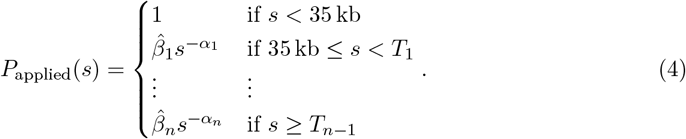

The transition points *T*_*i*_ are determined by the intersections of the local regressions in log-log space and verified by bootstrapping. The scaling constants 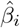 are recalculated for to ensure continuity at each boundary and to normalize the function such that *P*_applied_(35 kb) = 1. The distance-corrected interaction scores are generated by multiplying the raw co-accessibility scores or DL predictions by the corresponding value of *P*_applied_(*s*).

To evaluate the GMM-based piecewise model, we benchmarked its performance against a global power-law model with a standard exponential ansatz *P* (*s*) = *e*^−*λs*^. Each penalty function was compared against raw Hi-C loop profiles for every individual dataset. We quantified model accuracy using *R*^2^ scores in log-log space to assess fit quality across the 5 Mb range. Detailed performance metrics are provided in the Supplementary Information (Supplementary Table S2 and Figure S3).

### 2.4 Calculation of Deep Learning-based interaction scores

Deep learning (DL)-based interaction scores for maize were generated using `GenomicLinks`, a sequence-based model that predicts chromatin looping potential directly from DNA [21]. The `GenomicLinks` architecture utilizes a combination of convolutional neural networks (CNNs) and long short-term memory (LSTM) layers, previously trained on experimental HiChIP data (Maize leaf tissue, targeting H3K4me3 and H3K27me3) [31]. We applied the pre-trained model to all pairwise combinations of the HiChIP anchor regions (2,500 bp length) across the maize genome. The model generates a raw DL score for each pair, representing the predicted interaction potential. To ensure a direct comparison with our other metrics, these predictions were filtered to a genomic distance range of 0 to 500 kb and corrected using the species-specific penalty function as described above.

### 2.5 Polymer model simulations

To characterize the relationship between the distance-decay exponent *α* and resulting chromatin topologies, we performed coarse-grained Monte Carlo simulations using the Pivot Algorithm [40]. Chains of *N* = 10, 000 steps were simulated on a cubic lattice across four canonical folding regimes: Random Walk (RW, *α* ≈ 1.5), Self-Avoiding Walk (SAW, *α* ≈ 1.76), and both models under spherical confinement to approximate Fractal Globule-like scaling (*α* ≈ 1.0). Ensembles were equilibrated using a number of pivot moves proportional to the chain length to ensure efficient sampling of the conformational space. The scaling exponent *α* for each model was empirically derived via linear regression of log_10_(*P* (*s*)) versus log_10_(*s*) for pairwise genomic separations *s >* 10 steps. Visualizations of the 3D conformations and corresponding *P* (*s*) decay curves were generated to validate the physical basis of the scaling exponents (see Supplementary Fig. S4 and Figure 1E).

## 3 Results and Discussion

### 3.1 Multi-component power-law decay of Hi-C contacts reveals a multi-scale plant chromatin architecture

We analyzed Hi-C contact–distance decay profiles from soybean, rice, and maize across multiple tissues to quantify how contact probability changes with genomic separation. On log–log axes, Hi-C contact frequency versus distance was consistently piecewise linear, indicating that a single power-law exponent is insufficient to describe chromatin contact scaling across distances [41] (Figure 2). We therefore fit each profile with a Gaussian mixture model (GMM) to identify multiple scaling regimes and estimate a set of decay exponents {*α*_*i*_} and their transition points (Figure 2; Figure 3A, E, I; Methods). Tissue-specific profiles typically required two to four regimes in soybean and rice but only one or two in maize. For global consensus models (pooled across tissues within species), the GMM identified three regimes in soybean and two regimes in rice and maize (Supplementary Table S1).

**Figure 2.**
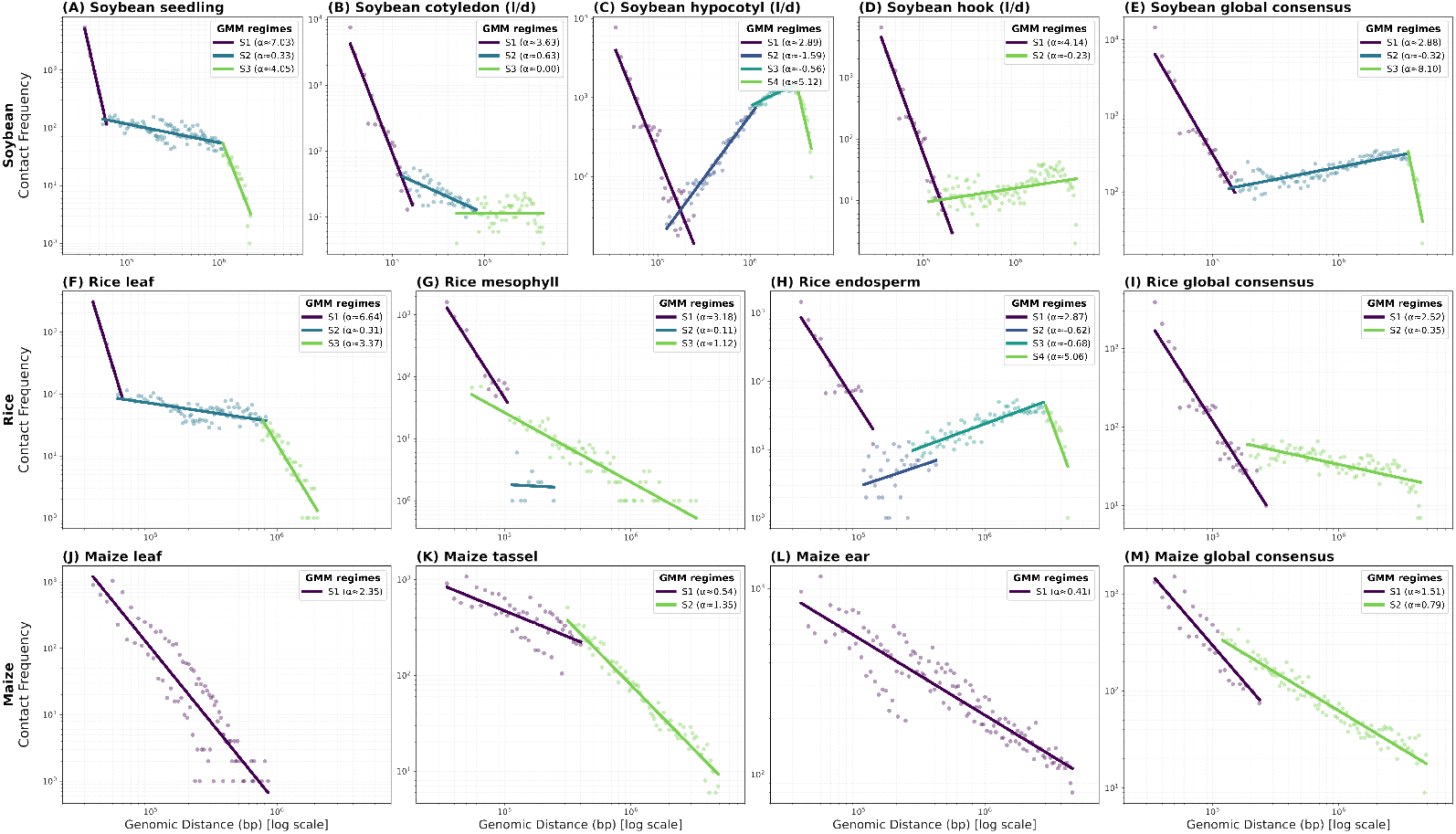
Tissue-specific GMM decomposition of chromatin interaction profiles. Hi-C contact frequencies (points) show that chromatin scaling is piecewise linear in log–log space, requiring multi-component functions to capture distinct architectural regimes. Gaussian Mixture Modeling (GMM) identifies these interaction subpopulations (*S*_1_ to *S*_4_), with each segment representing a unique power-law decay exponent derived from cluster covariance. These profiles demonstrate that interaction scaling is both species- and tissue-specific. Ten independent datasets are shown: soybean (**A–D**: seedling, cotyledon, hypocotyl, hook), rice (**F–H**: leaf, mesophyll, endosperm), and maize (**J–L**: leaf, tassel, ear), alongside their respective consensus profiles (**E, I, M**).

**Figure 3.**
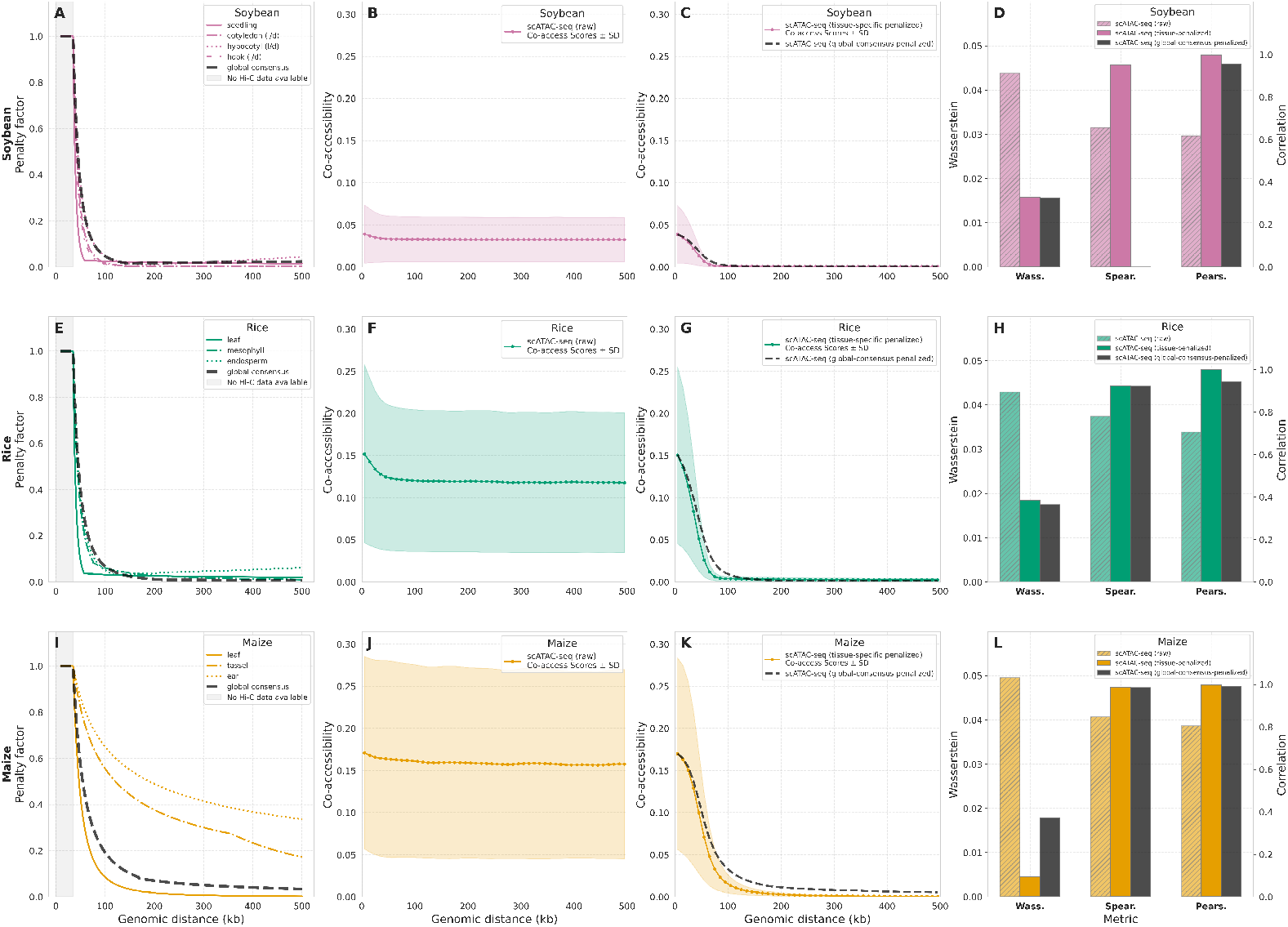
Application and statistical validation of species-specific and global consensus penalty functions. **(A, E, I)** Species-specific and global consensus penalty functions derived via GMM decomposition of Hi-C interaction counts. Hi-C data constraints limit the penalty domain to distances *>* 35 kb. **(B, F, J)** Raw co-accessibility scores show minimal decay with increasing genomic distance, overestimating distal contacts. **(C, G, K)** Application of tissue-specific (colored line) and global consensus (black dashed line) penalties. Both models successfully transform raw scores into penalized profiles that exhibit Hi-C-like decay. Shaded areas represent the standard deviation of co-accessibility scores across ACR pairs. **(D, H, L)** Statistical validation for soybean, rice, and maize. Compared to raw scores (hatched bars), penalized scores from tissue-specific (colored bars) and global consensus (gray bars) models show markedly improved concordance with Hi-C data, evidenced by lower Wasserstein distances and higher Pearson correlation coefficients.

The inferred regimes align with known architectural scales. In rice and soybean, the first regime extended to bootstrapped transition points *T*_1_ with medians of 120.4 kb (rice) and 140.2 kb (soybean) and showed steep decay (*α* ≈ 2.6–3.0), consistent with strongly partitioned short-range organization (Figure 1E). These *T*_1_ thresholds coincide with independent loop annotations, encompassing 97% of identified loops in rice [42] and 70% in soybean [28].

Beyond *T*_1_, both rice and soybean entered a second regime with weak or near-flat scaling (*α* = 0.37 in rice; *α* = −0.33 in soybean), consistent with mid-range enrichment and domain-scale organization. A strictly monotonic distance penalty would therefore over-suppress enriched distance ranges, motivating a multi-regime penalty design. In contrast, maize exhibited a more monotonic scaling behavior. While the global consensus model supported a transition at ∼168 kb, several tissues were best captured by a single power-law (e.g., leaf, *α* = 2.35; Figure 2).

### 3.2 Co-accessibility scores and deep-learning predictions overestimate long-range interactions

To quantify distance bias in scalable interaction proxies, we re-analyzed leaf scATAC-seq datasets from soybean [18], rice [29], and maize [27] and compared their co-accessibility distance profiles against tissue-matched Hi-C (leaf for rice and maize; seedling for soybean). Co-accessibility inference from scATAC-seq is challenged by sparsity, so we followed standard preprocessing by constructing pseudocells using a data-driven *k*-nearest-neighbors strategy [27]. We then computed pairwise correlations of normalized accessibility across pseudocells for ACR pairs separated by 2 to 500 kb. This yielded 6.3 million positively co-accessible ACR pairs in soybean, 3.8 million in rice, and 3.4 million in maize.

A side-by-side comparison with Hi-C revealed a consistent distortion: raw co-accessibility scores show little to no decay with increasing genomic distance, with elevated interaction scores at large separations relative to Hi-C (Figure 1B–D; Figure 3B, F, J). Uncorrected co-accessibility exhibited very high long-range false positive rates (FPR), exceeding 0.90 across datasets (Supplementary Table S2).

Distance bias is not specific to co-accessibility. We observed the same pattern in deep-learning (DL) interaction proxies, which often omit explicit genomic distance as a feature during training [43, 16, 21]. Using `GenomicLinks` predictions for maize [21], raw DL scores were largely distance-independent across the 0–500 kb interval (Figure 4B), mirroring co-accessibility inflation and overestimating distal interactions relative to the Hi-C reference. Together, these analyses motivate a correction that is distance-aware, learned from empirical Hi-C decay in the relevant species, and portable across proxy score types.

**Figure 4.**
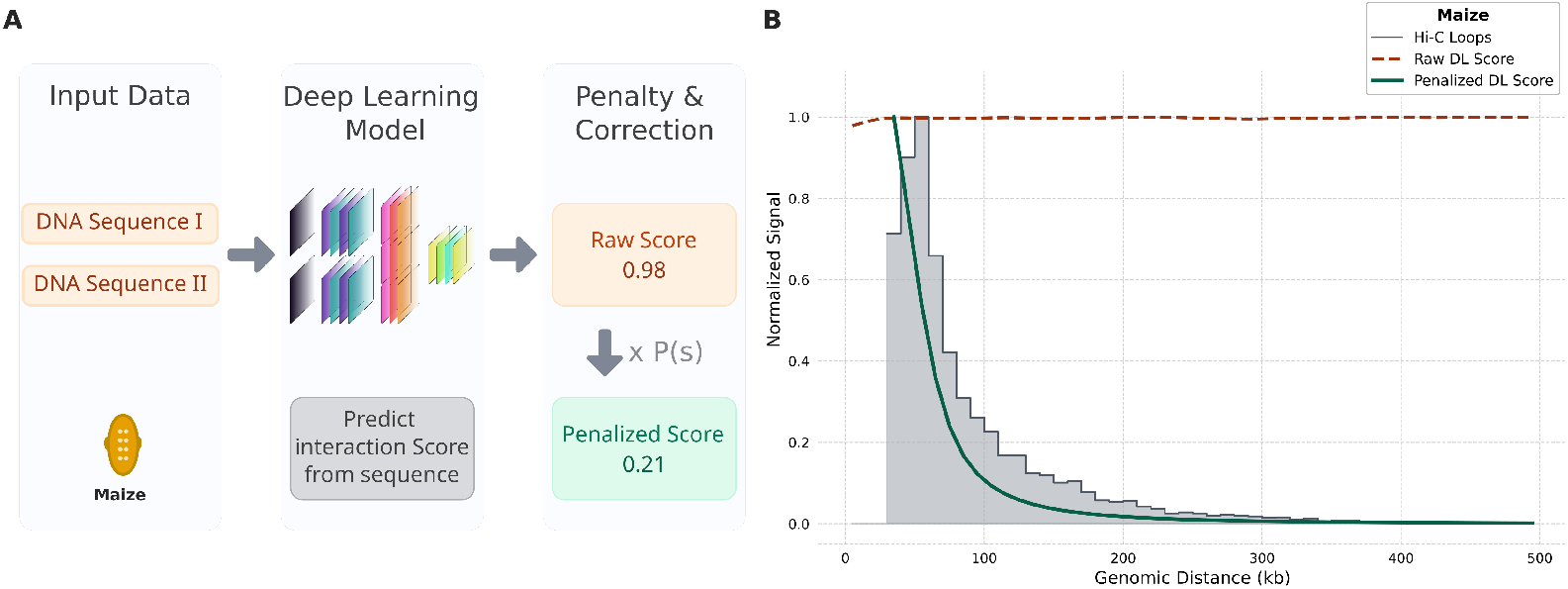
A polymer-based penalty corrects distance bias in Deep Learning predictions for maize. **(A)** Workflow for correcting Deep Learning (DL) scores. The GenomicLinks model predicts raw interaction scores from DNA sequences, which are subsequently corrected using the species-specific penalty function derived from Hi-C data. **(B)** Distance-bias correction in maize. Raw DL scores (dark red dashed line) are mostly independent of genomic distance, overestimating long-range contacts relative to the Hi-C reference (grey histogram). Applying the polymer-based penalty produces a corrected profile (dark green solid line) that closely recapitulates the experimental Hi-C decay. All profiles are normalized to 1.0.

### 3.3 A GMM-based penalty corrects distance bias in co-accessibility and deep-learning predictions

Several methods implement distance-related adjustments for co-accessibility, but many rely on fixed exponents, heuristic cutoffs, or non-transferable background assumptions [44, 45, 17]. In plants, both decay exponents and regime boundaries vary across species and tissues, making universal one-parameter corrections suboptimal. We therefore derived penalties directly from GMM-fitted Hi-C decay profiles. Across the 35–500 kb interval used for scATAC-seq bench-marking, the multi-regime model outperformed a single-exponential fit, with at least an 11.1-fold reduction in residual sum of squares (RSS) across all datasets (Supplementary Fig. S5).

Applying tissue-specific penalties to raw co-accessibility scores reshaped distance-independent profiles into Hi-C-like decay curves (Figure 1B–D; Figure 3C, G, K). Quantitatively, corrected profiles showed improved concordance with Hi-C (higher Pearson, lower Wasserstein) and markedly improved specificity. FPR dropped from above 90% in raw profiles to 2.2% in maize, 39.6% in rice, and 33.4% in soybean, while average F1 increased from 0.15 to 0.83 across tissue-specific models (Supplementary Table S2). The soybean global consensus revealed a statistical uniqueness: while it achieved a strong F1 score (0.814), the Spearman correlation dropped to −0.014. This is localized to a small genomic segment where the penalty slightly upregulates co-accessibility scores due to a non-monotonic scaling regime (*α <* 0), creating a rank-order disagreement (Supplementary Fig. S6). Global consensus penalties provided a robust default when tissue-matched Hi-C was unavailable. In maize and rice, global consensus performance was close to tissue-specific models across profile and error-rate metrics (Supplementary Table S2). Across species, long-range false positives were reduced by an average of 73% for tissue-specific penalties and 66% for global consensus penalties.

To test biological plausibility, we stratified maize co-accessible ACR pairs by penalty magnitude (∆ co-accessibility). The most penalized tertile was enriched for genic–genic and distal–distal links, while distal–proximal links were enriched among least penalized pairs (Figure 5A). Genic– genic pairs in the most penalized tertile also showed significantly higher co-expression values (Kruskal-Wallis test, *P <* 2.2 × 10^−16^; Figure 5B), consistent with the depletion of co-regulatory correlations that can inflate long-range proxy scores. At the same time, known long-range cis-regulatory loci associated with *ZmRAP2.7* [46] and *bx1* [47] retained elevated penalized co-accessibility relative to background (Wilcoxon test, *P <* 5.9 × 10^−52^; Figure 5C–D). For example, *Vgt1* and *Vgt1.2* [48] regulatory loci exhibit penalized signal above background with *ZmRAP2.7*, though *Vgt1* is down-weighted more than *Vgt1.2* (Figure 5D).

**Figure 5.**
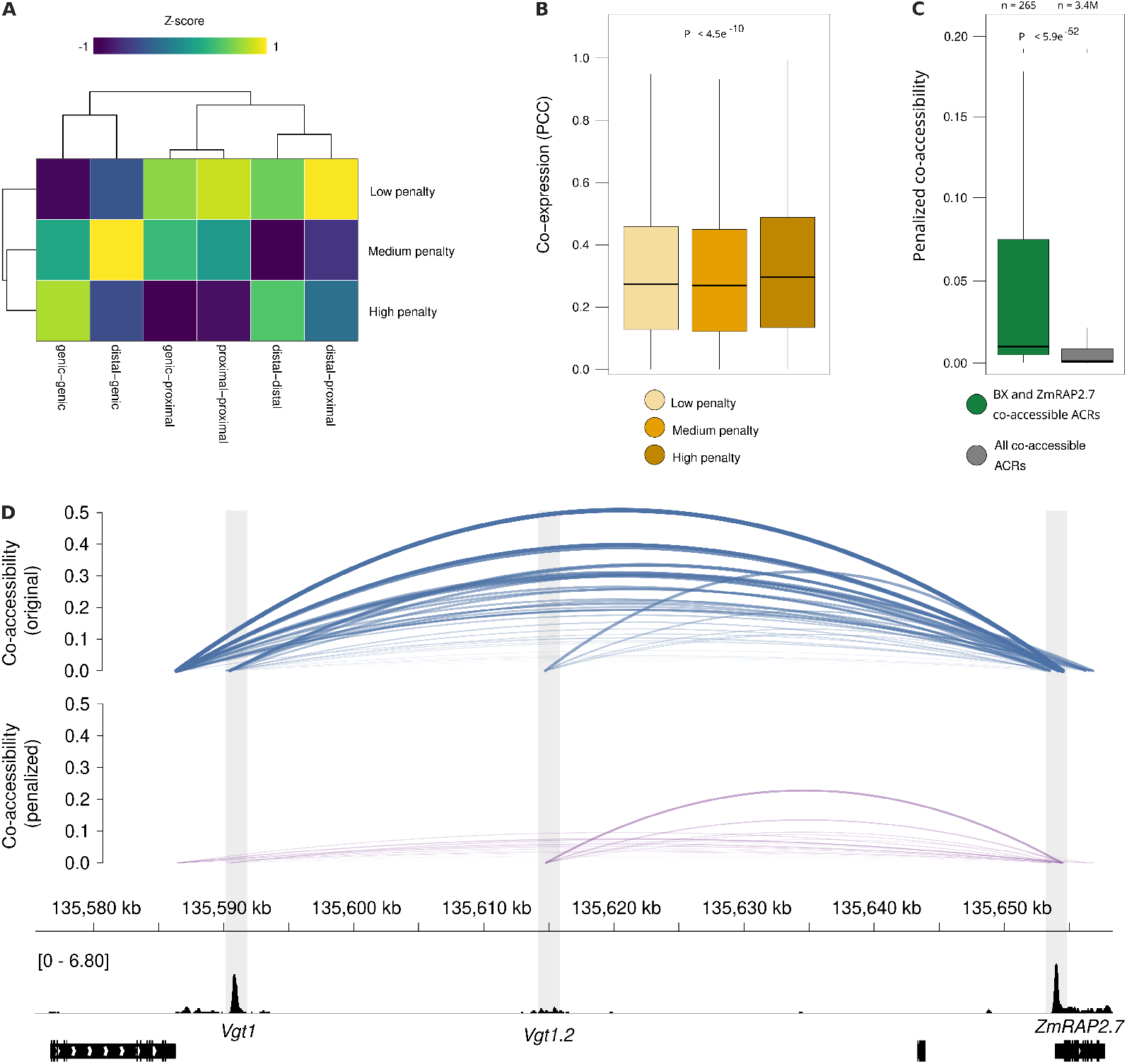
Penalty enriches for functional interactions and separates co-regulation from physical proximity in maize. **(A)** Heatmap of co-accessible ACR classification enrichment (*Z*-score) in low, medium, and high penalized tertiles. Proximal–distal links are enriched in the low penalty group, while genic–genic links are most penalized. **(B)** Boxplots showing that gene pairs with the highest penalty scores also have the highest co-expression (PCC, Pearson Correlation Coefficient), suggesting the penalty effectively down-weights co-regulated, non-interacting loci. **(C)** Known functional long-range interactions (at the *BX1* and *ZmRAP2.7* loci) retain significantly higher penalized co-accessibility scores compared to the global set of all co-accessible ACRs. **(D)** Genome browser view of raw (top) and penalized (middle) co-accessibility scores, and chromatin accessibility profiles (bottom) at the *Vgt1*–*ZmRAP2.7* locus. *Vgt1*, Vegetative to generative transition 1. *ZmRAP2.7, Zea mays* RELATED TO APETALA2.

Finally, applying the maize species-specific penalty to `GenomicLinks` predictions converted distance-independent raw DL profiles into curves aligned with Hi-C decay (Figure 4B), confirming the method functions as a modular and broadly applicable tool for correcting distance bias in diverse interaction proxies.

## 4 Conclusion

We introduced a distance-aware correction framework for scalable chromatin-interaction proxies that systematically inflate long-range contacts. By learning multi-regime distance-decay behavior from Hi-C using Gaussian mixture modeling, we derive tissue-specific and global consensus penalty functions and apply them to co-accessibility and deep-learning interaction scores. This correction restores Hi-C-like distance profiles and reduces long-range false positives by about 70% on average across maize, rice, and soybean, while preserving biologically meaningful interaction signal.

While this framework provides an effective correction, it currently relies on bulk Hi-C data for calibration. However, our global consensus penalties perform nearly as well as tissue-specific models, achieving Spearman correlations as high as 0.98 in maize and 0.92 in rice. Key future directions include integrating these parameters into dynamic polymer models [49] and incorporating distance awareness directly into the deep-learning training process. We release code and fitted parameters (jlab-github) to support adoption and extension to additional plant species and tissues.

## Declarations

### Ethics approval and consent to participate

Not applicable.

### Consent for publication

Not applicable.

### Availability of data and materials

The open-source code and derived parameters generated during this study are publicly available on GitHub (https://github.com/jlab-code/polymer-penalty). The processed datasets are available through Zenodo (https://doi.org/10.5281/zenodo.18607660). All Hi-C and scATAC-seq datasets re-analyzed in this study are publicly available; a full list of original sources and accession numbers is provided in Supplementary Table S3.

## Authors’ contributions

LS performed the data analysis, contributed to the study design, and wrote the manuscript. FGC provided essential data and contributed to the analysis pipeline. APM and FJ supervised the project, conceived the study, and edited the manuscript. All authors read and approved the final manuscript.

## Acknowledgements

We thank members of the Johannes lab for helpful feedback.

## References

[1] Boyan Bonev and Giacomo Cavalli. “Organization and function of the 3D genome”. en. In: Nat Rev Genet 17.11 (Oct. 2016), pp. 661–678.

[2] M Jordan Rowley and Victor G Corces. “Organizational principles of 3D genome architecture”. en. In: Nat Rev Genet 19.12 (Oct. 2018), pp. 789–800.

[3] Katherine Domb, Nan Wang, Guillaume Hummel, and Chang Liu. “Spatial Features and Functional Implications of Plant 3D Genome Organization”. en. In: Annu Rev Plant Biol 73 (May 2022), pp. 173–200.

[4] Suhas S P Rao, Miriam H Huntley, Neva C Durand, Elena K Stamenova, Ivan D Bochkov, James T Robinson, Adrian L Sanborn, Ido Machol, Arina D Omer, Eric S Lander, et al. “A 3D map of the human genome at kilobase resolution reveals principles of chromatin looping”. en. In: Cell 159.7 (Dec. 2014), pp. 1665–1680.

[5] Delfina Gagliardi and Pablo A Manavella. “Short-range regulatory chromatin loops in plants”. en. In: New Phytol 228.2 (Oct. 2020), pp. 466–471.

[6] Anil Panigrahi and Bert W O’Malley. “Mechanisms of enhancer action: the known and the unknown”. en. In: Genome Biol 22.1 (Apr. 2021), pp. 1–30.

[7] Jesse R Dixon, Siddarth Selvaraj, Feng Yue, Audrey Kim, Yan Li, Yin Shen, Ming Hu, Jun S Liu, and Bing Ren. “Topological Domains in Mammalian Genomes Identified by Analysis of Chromatin Interactions”. In: Nature 485.7398 (Apr. 2012), p. 376.

[8] Jonathan A Beagan and Jennifer E Phillips-Cremins. “On the existence and functionality of topologically associating domains”. en. In: Nat Genet 52.1 (Jan. 2020), pp. 8–16.

[9] Quentin Szabo, Frédéric Bantignies, and Giacomo Cavalli. “Principles of genome folding into topologically associating domains”. In: Sci Adv 5.4 (Apr. 2019), eaaw1668.

[10] E Lieberman-Aiden, N L van Berkum, L Williams, M Imakaev, T Ragoczy, A Telling, I Amit, B R Lajoie, P J Sabo, M O Dorschner, et al. “Comprehensive mapping of long-range interactions reveals folding principles of the human genome”. In: Science 326.5950 (Oct. 2009), pp. 289–296.

[11] Hannah L Harris, Huiya Gu, Moshe Olshansky, Ailun Wang, Irene Farabella, Yossi Eliaz, Achyuth Kalluchi, Akshay Krishna, Mozes Jacobs, Gesine Cauer, et al. “Chromatin alternates between A and B compartments at kilobase scale for subgenic organization”. en. In: Nat Commun 14.1 (June 2023), pp. 1–17.

[12] Pengfei Dong, Xiaoyu Tu, Haoxuan Li, Jianhua Zhang, Donald Grierson, Pinghua Li, and Silin Zhong. “3D Chromatin Architecture of Large Plant Genomes Determined by Local A/B Compartments”. In: Mol Plant 10.12 (Dec. 2017), pp. 1497–1509.

[13] Pengfei Dong, Xiaoyu Tu, Haoxuan Li, Jianhua Zhang, Donald Grierson, Pinghua Li, and Silin Zhong. “Tissue-specific Hi-C analyses of rice, foxtail millet and maize suggest non-canonical function of plant chromatin domains”. en. In: J Integr Plant Biol 62.2 (Feb. 2020), pp. 201–217.

[14] Takashi Nagano, Yaniv Lubling, Eitan Yaffe, Steven W Wingett, Wendy Dean, Amos Tanay, and Peter Fraser. “Single-cell Hi-C for genome-wide detection of chromatin interactions that occur simultaneously in a single cell”. en. In: Nat Protoc 10.12 (Nov. 2015), pp. 1986–2003.

[15] Alexandre P Marand, Zongliang Chen, Andrea Gallavotti, and Robert J Schmitz. “A cis-regulatory atlas in maize at single-cell resolution”. en. In: Cell 184.11 (May 2021), 3041–3055.e21.

[16] Ron Schwessinger, Matthew Gosden, Damien Downes, Richard C Brown, A Marieke Oudelaar, Jelena Telenius, Yee Whye Teh, Gerton Lunter, and Jim R Hughes. “DeepC: predicting 3D genome folding using megabase-scale transfer learning”. en. In: Nat Methods 17.11 (Nov. 2020), pp. 1118–1124.

[17] Hannah A Pliner, Jonathan S Packer, José L McFaline-Figueroa, Darren A Cusanovich, Riza M Daza, Delasa Aghamirzaie, Sanjay Srivatsan, Xiaojie Qiu, Dana Jackson, Anna Minkina, et al. “Cicero Predicts cis-Regulatory DNA Interactions from Single-Cell Chromatin Accessibility Data”. en. In: Mol Cell 71.5 (Sept. 2018), 858–871.e8.

[18] Xuan Zhang, Ziliang Luo, Alexandre P Marand, Haidong Yan, Hosung Jang, Sohyun Bang, John P Mendieta, Mark A A Minow, and Robert J Schmitz. “A spatially resolved multi-omic single-cell atlas of soybean development”. en. In: Cell 188.2 (Jan. 2025), 550–567.e19.

[19] Robert S Piecyk, Luca Schlegel, and Frank Johannes. “Predicting 3D chromatin interactions from DNA sequence using Deep Learning”. en. In: Comput Struct Biotechnol J 20 (June 2022), pp. 3439–3448.

[20] James Zou, Mikael Huss, Abubakar Abid, Pejman Mohammadi, Ali Torkamani, and Amalio Telenti. “A primer on deep learning in genomics”. en. In: Nat Genet 51.1 (Jan. 2019), pp. 12–18.

[21] Luca Schlegel, Rohan Bhardwaj, Yadollah Shahryary, Defne Demirtürk, Alexandre P Marand, Robert J Schmitz, and Frank Johannes. “GenomicLinks: deep learning predictions of 3D chromatin interactions in the maize genome”. en. In: NAR Genom Bioinform 6.3 (Sept. 2024), lqae123.

[22] Benjamin P. G. Wall, Mary Nguyen, J. Chuck Harrell, and Mikhail G. Dozmorov. “Machine and Deep Learning Methods for Predicting 3D Genome Organization”. In: Methods in Molecular Biology 2856 (2025), pp. 357–400.

[23] H. Chen, C. Lareau, T. Andreani, M. E. Vinyard, A. J. Moore, B. Cavaza, K. Fejtova, Z. Shao, A. Augello, R. Misra, et al. “Assessment of computational methods for the analysis of single-cell ATAC-seq data”. In: Genome Biology 20.1 (2019), p. 241.

[24] Dong-Sung Lee, Chongyuan Luo, Jingtian Zhou, Sahaana Chandran, Angeline Rivkin, Anna Bartlett, Joseph R. Nery, Conor Fitzpatrick, Carolyn O’Connor, Jesse R. Dixon, et al. “Simultaneous profiling of 3D genome structure and DNA methylation in single human cells”. In: Nature Methods 16.10 (2019), pp. 999–1006.

[25] Jixian Zhai, Dong-Hoon Jeong, Emanuele De Paoli, Sunhee Park, Benjamin D Rosen, Yupeng Li, Alvaro J González, Zhe Yan, Sherry L Kitto, Michael A Grusak, et al. “MicroRNAs as master regulators of the plant NB-LRR defense gene family via the production of phased, trans-acting siRNAs”. en. In: Genes Dev 25.23 (Dec. 2011), pp. 2540–2553.

[26] Yunlong Wang, Siyuan Kong, Cong Zhou, Yanfang Wang, Yubo Zhang, Yaping Fang, and Guoliang Li. “A review of deep learning models for the prediction of chromatin interactions with DNA and epigenomic profiles”. en. In: Brief Bioinform 26.1 (Dec. 2024), bbae651.

[27] Alexandre P Marand, Luguang Jiang, Fabio Gomez-Cano, Mark A A Minow, Xuan Zhang, John P Mendieta, Ziliang Luo, Sohyun Bang, Haidong Yan, Cullan Meyer, et al. “The genetic architecture of cell type-specific cis regulation in maize”. en. In: Science 388.6744 (Apr. 2025), eads6601.

[28] Longfei Wang, Guanghong Jia, Xinyu Jiang, Shuai Cao, Z Jeffrey Chen, and Qingxin Song. “Altered chromatin architecture and gene expression during polyploidization and domestication of soybean”. en. In: Plant Cell 33.5 (Mar. 2021), pp. 1430–1446.

[29] Haidong Yan, John P Mendieta, Xuan Zhang, Alexandre P Marand, Yan Liang, Ziliang Luo, Thomas Roulé, Doris Wagner, Xiaoyu Tu, Yonghong Wang, et al. “Evolution of cell-type-specific accessible chromatin regions and the cis-regulatory elements that drive lineage-specific innovation”. In: bioRxiv (2024).

[30] Chang Liu, Ying-Juan Cheng, Jia-Wei Wang, and Detlef Weigel. “Prominent topologically associated domains differentiate global chromatin packing in rice from Arabidopsis”. en. In: Nat Plants 3.9 (Sept. 2017), pp. 742–748.

[31] William A Ricci, Zefu Lu, Lexiang Ji, Alexandre P Marand, Christina L Ethridge, Nathalie G Murphy, Jaclyn M Noshay, Mary Galli, María Katherine Mejía-Guerra, Maria Colomé-Tatché, et al. “Widespread long-range cis-regulatory elements in the maize genome”. en. In: Nat Plants 5.12 (Dec. 2019), pp. 1237–1249.

[32] Zhu Li, Linhua Sun, Xiao Xu, Yutong Liu, Hang He, and Xing Wang Deng. “Light control of three-dimensional chromatin organization in soybean”. en. In: Plant Biotechnol J 22.9 (Sept. 2024), pp. 2596–2611.

[33] Nicolas Servant, Nelle Varoquaux, Bryan R Lajoie, Eric Viara, Chong-Jian Chen, Jean-Philippe Vert, Edith Heard, Job Dekker, and Emmanuel Barillot. “HiC-Pro: an optimized and flexible pipeline for Hi-C data processing”. en. In: Genome Biol 16 (Dec. 2015), p. 259.

[34] Neva C Durand, Muhammad S Shamim, Ido Machol, Suhas S P Rao, Miriam H Huntley, Eric S Lander, and Erez Lieberman Aiden. “Juicer Provides a One-Click System for Analyzing Loop-Resolution Hi-C Experiments”. en. In: Cell Syst 3.1 (July 2016), pp. 95–98.

[35] Scott C Stelpflug, Rajandeep S Sekhon, Brieanne Vaillancourt, Candice N Hirsch, C Robin Buell, Natalia de Leon, and Shawn M Kaeppler. “An Expanded Maize Gene Expression Atlas based on RNA Sequencing and its Use to Explore Root Development”. In: Plant Genome 9.1 (Mar. 2016), lantgenome2015.04.0025.

[36] Statistical Physics of Macromolecules. en. Amer Inst of Physics, 1994.

[37] Leonid A Mirny. “The fractal globule as a model of chromatin architecture in the cell”. en. In: Chromosome Res 19.1 (Jan. 2011), pp. 37–51.

[38] F. Pedregosa, G. Varoquaux, A. Gramfort, V. Michel, B. Thirion, O. Grisel, M. Blondel, P. Prettenhofer, R. Weiss, V. Dubourg, et al. “Scikit-learn: Machine Learning in Python”. In: Journal of Machine Learning Research 12 (2011), pp. 2825–2830.

[39] Robert E. Kass and Adrian E. Raftery. “Bayes Factors”. In: Journal of the American Statistical Association 90.430 (1995), pp. 773–795.

[40] Neal Madras and Alan D Sokal. “The pivot algorithm: A highly e?icient Monte Carlo method for the self-avoiding walk”. en. In: J. Stat. Phys. 50. 1-2 (Jan. 1988), pp. 109–186.

[41] Adrian L Sanborn, Suhas S P Rao, Su-Chen Huang, Neva C Durand, Miriam H Huntley, Andrew I Jewett, Ivan D Bochkov, Dharmaraj Chinnappan, Ashok Cutkosky, Jian Li, et al. “Chromatin extrusion explains key features of loop and domain formation in wild-type and engineered genomes”. en. In: Proc Natl Acad Sci U S A 112.47 (Nov. 2015), E6456–65.

[42] Lun Zhao, Shuangqi Wang, Zhilin Cao, Weizhi Ouyang, Qing Zhang, Liang Xie, Ruiqin Zheng, Minrong Guo, Meng Ma, Zhe Hu, et al. “Chromatin loops associated with active genes and heterochromatin shape rice genome architecture for transcriptional regulation”. en. In: Nat Commun 10.1 (Aug. 2019), p. 3640.

[43] Geoff Fudenberg, David R Kelley, and Katherine S Pollard. “Predicting 3D genome folding from DNA sequence with Akita”. en. In: Nat Methods 17.11 (Nov. 2020), pp. 1111–1117.

[44] Chenfei Wang, Dongqing Sun, Xin Huang, Changxin Wan, Ziyi Li, Ya Han, Qian Qin, Jingyu Fan, Xintao Qiu, Yingtian Xie, et al. “Integrative analyses of single-cell transcriptome and regulome using MAESTRO”. en. In: Genome Biology 21.1 (Aug. 2020), p. 198.

[45] Jeffrey M Granja, M Ryan Corces, Sarah E Pierce, S Tansu Bagdatli, Hani Choudhry, Howard Y Chang, and William J Greenleaf. “ArchR is a scalable software package for integrative single-cell chromatin accessibility analysis”. en. In: Nat Genet 53.3 (Mar. 2021), pp. 403–411.

[46] Silvio Salvi, Giorgio Sponza, Michele Morgante, Dwight Tomes, Xiaomu Niu, Kevin A Fengler, Robert Meeley, Evgueni V Ananiev, Sergei Svitashev, Edward Bruggemann, et al. “Conserved noncoding genomic sequences associated with a flowering-time quantitative trait locus in maize”. en. In: Proc Natl Acad Sci U S A 104.27 (July 2007), pp. 11376–11381.

[47] Linlin Zheng, Michael D McMullen, Eva Bauer, Chris-Carolin Schön, Alfons Gierl, and Monika Frey. “Prolonged expression of the BX1 signature enzyme is associated with a recombination hotspot in the benzoxazinoid gene cluster in Zea mays”. en. In: J Exp Bot 66.13 (July 2015), pp. 3917–3930.

[48] Julia Engelhorn, Samantha J Snodgrass, Amelie Kok, Arun S Seetharam, Michael Schneider, Tatjana Kiwit, Ayush Singh, Michael Banf, Duong Thi Hai Doan, Merritt Khaipho-Burch, et al. “Genetic variation at transcription factor binding sites largely explains phenotypic heritability in maize”. en. In: Nat Genet (Aug. 2025), pp. 1–10.

[49] Vinayak Vinayak, Ramin Basir, Rosela Golloshi, Joshua Toth, Lucas Sant’Anna, Melike Lakadamyali, Rachel Patton McCord, and Vivek B Shenoy. “Polymer model integrates imaging and sequencing to reveal how nanoscale heterochromatin domains influence gene expression”. en. In: Nat Commun 16.1 (Apr. 2025), p. 3816.

