## Supplementary Information for "A multi-component power-law penalty corrects distance bias in single-cell co-accessibility and deep-learning chromatin interaction predictions"

#### Supplemental Tables

| DATASET | K (REGIMES) | COMPONENT (I) | $\alpha_i$ (DECAY) | $\pi_i$ (WEIGHT) | $\beta_i$ (GMM SCALING) | $\hat{\beta}_i$ (APPLIED SCALING) | TRANSITIONS (KB) |
|---|---|---|---|---|---|---|---|
| Soybean seedling | 3 | 1 | 7.03 | 0.04 | $4.66 \times 10^{35}$ | $8.62 \times 10^{31}$ | 57.4, 1087.4 |
| | | 2 | 0.32 | 0.75 | $5.20 \times 10^{3}$ | $9.53 \times 10^{-1}$ | |
| | | 3 | 4.13 | 0.22 | $5.22 \times 10^{26}$ | $3.10 \times 10^{22}$ | |
| Soybean cotyledon (l/d) | 3 | 1 | 3.76 | 0.17 | $5.61 \times 10^{20}$ | $3.05 \times 10^{16}$ | 146.5, 636.4 |
| | | 2 | 0.64 | 0.42 | $4.32 \times 10^{4}$ | $1.61 \times 10^{1}$ | |
| | | 3 | -0.03 | 0.41 | $5.85 \times 10^{0}$ | $2.70 \times 10^{-3}$ | |
| Soybean hypocotyl (l/d) | 4 | 1 | 2.90 | 0.24 | $6.40 \times 10^{16}$ | $1.34 \times 10^{13}$ | 175.0, 1099.4, 3069.6 |
| | | 2 | -1.58 | 0.43 | $1.88 \times 10^{-7}$ | $4.66 \times 10^{-11}$ | |
| | | 3 | -0.56 | 0.22 | $2.76 \times 10^{-1}$ | $8.44 \times 10^{-5}$ | |
| | | 4 | 5.25 | 0.10 | $1.45 \times 10^{37}$ | $7.18 \times 10^{32}$ | |
| Soybean hook (l/d) | 2 | 1 | 4.14 | 0.18 | $2.85 \times 10^{22}$ | $6.74 \times 10^{18}$ | 153.5 |
| | | 2 | -0.23 | 0.82 | $6.21 \times 10^{-1}$ | $1.38 \times 10^{-4}$ | |
| Soybean global consensus | 3 | 1 | 2.91 | 0.17 | $1.14 \times 10^{17}$ | $1.19 \times 10^{13}$ | 142.3, 3484.6 |
| | | 2 | -0.33 | 0.76 | $2.12 \times 10^{0}$ | $3.74 \times 10^{-4}$ | |
| | | 3 | 8.13 | 0.07 | $5.38 \times 10^{55}$ | $6.21 \times 10^{51}$ | |
| Rice leaf | 3 | 1 | 6.65 | 0.04 | $5.05 \times 10^{33}$ | $1.42 \times 10^{30}$ | 57.4, 792.2 |
| | | 2 | 0.28 | 0.65 | $2.40 \times 10^{3}$ | $8.18 \times 10^{-1}$ | |
| | | 3 | 3.45 | 0.31 | $1.18 \times 10^{22}$ | $8.32 \times 10^{17}$ | |
| Rice mesophyll | 3 | 1 | 3.15 | 0.15 | $2.70 \times 10^{17}$ | $2.96 \times 10^{14}$ | 76.0, 614.2 |
| | | 2 | 1.12 | 0.76 | $3.17 \times 10^{7}$ | $4.93 \times 10^{-3}$ | |
| | | 3 | 0.33 | 0.09 | $9.27 \times 10^{2}$ | $8.39 \times 10^{3}$ | |
| Rice endosperm | 4 | 1 | 2.84 | 0.13 | $6.80 \times 10^{15}$ | $1.06 \times 10^{13}$ | 119.6, 332.7, 2964.5 |
| | | 2 | -0.47 | 0.21 | $1.04 \times 10^{-1}$ | $2.71 \times 10^{-6}$ | |
| | | 3 | -0.66 | 0.55 | $9.00 \times 10^{-3}$ | $2.29 \times 10^{-6}$ | |
| | | 4 | 5.51 | 0.11 | $7.76 \times 10^{37}$ | $3.16 \times 10^{31}$ | |
| Rice global consensus | 2 | 1 | 2.56 | 0.25 | $7.05 \times 10^{14}$ | $2.80 \times 10^{11}$ | 224.4 |
| | | 2 | 0.36 | 0.75 | $1.23 \times 10^{3}$ | $2.64 \times 10^{0}$ | |
| Maize leaf | 1 | 1 | 2.35 | 1.00 | $5.69 \times 10^{13}$ | $4.71 \times 10^{10}$ | — |
| Maize tassel | 2 | 1 | 0.56 | 0.42 | $3.12 \times 10^{5}$ | $2.87 \times 10^{2}$ | 355.0 |
| | | 2 | 1.37 | 0.58 | $9.83 \times 10^{9}$ | $1.13 \times 10^{7}$ | |
| Maize ear | 1 | 1 | 0.41 | 1.00 | $6.36 \times 10^{4}$ | $7.61 \times 10^{1}$ | — |
| Maize global consensus | 2 | 1 | 1.57 | 0.19 | $2.09 \times 10^{10}$ | $7.18 \times 10^{6}$ | 167.9 |
| | | 2 | 0.79 | 0.81 | $1.65 \times 10^{6}$ | $2.29 \times 10^{3}$ | |

Table 1: **GMM-derived parameters for tissue-specific penalty functions.** This table provides the specific parameters used to construct the piecewise penalty function $P_{\text{applied}}(s)$. Parameters were derived by fitting a Gaussian Mixture Model (GMM) to the log–log distribution of Hi-C loop counts. The final correction follows a multi-regime power-law architecture: $P_{\text{applied}}(s) = \hat{\beta}_i s^{-\alpha_i}$ for genomic distances $s$ within identified regimes $[T_{i-1}, T_i]$. For each component $i$, $\alpha_i$ represents the decay exponent, $w_i$ the GMM weight, and $T_i$ the bootstrapped transition points. To maintain consistency with the numerical scale of co-accessibility scores (ranging from $-1.0$ to $1.0$), the resulting penalty is normalized such that $P_{\text{applied}}(35\text{ kb}) = 1.0$, ensuring no over-correction at the lower distance bound.

| SPECIES | MODEL | FPR | FNR | F1 SCORE | SPEARMAN | PEARSON | WASSERSTEIN |
|---|---|---|---|---|---|---|---|
| Soybean | Raw | 0.9308 | 0.0000 | 0.129 | 0.657 | 0.618 | 8.1605 |
| | Tissue-Spec | 0.3336 | 0.0000 | 0.800 | 0.952 | 0.999 | 2.9762 |
| | Global-Cons | 0.2999 | 0.0290 | 0.814 | -0.014 | 0.957 | 1.2686 |
| Rice | Raw | 0.9281 | 0.0000 | 0.134 | 0.779 | 0.706 | 8.0462 |
| | Tissue-Spec | 0.3961 | 0.0000 | 0.753 | 0.922 | 1.000 | 3.4761 |
| | Global-Cons | 0.2451 | 0.0843 | 0.828 | 0.922 | 0.943 | 1.4981 |
| Maize | Raw | 0.9005 | 0.0000 | 0.181 | 0.847 | 0.806 | 9.6565 |
| | Tissue-Spec | 0.0216 | 0.1073 | 0.934 | 0.986 | 0.997 | 0.3093 |
| | Global-Cons | 0.3834 | 0.0000 | 0.763 | 0.986 | 0.992 | 3.2945 |

Table 2: **Statistical and area-based validation of scATAC co-accessibility scores.** This table summarizes error rates and similarity metrics for raw and penalized scATAC-seq co-accessibility against Hi-C ground truth. False Positive Rate (FPR) and False Negative Rate (FNR) quantify the integrated areas of over- and under-prediction, respectively, normalized to the initial interaction frequency at 35 kb. The F1 Score represents the harmonic mean of area-based precision $(1-\text{FPR})$ and recall $(1-\text{FNR})$. Model performance is further assessed via Spearman ($\rho$) and Pearson ($r$) correlation coefficients and Wasserstein distance to quantify monotonic, linear, and distributional alignment.

| DATA | SPECIES | REPOSITORY | ACCESSION(S) | TISSUE DETAILS | PUBLICATION |
|---|---|---|---|---|---|
| Hi-C | Soybean | PRJNA | PRJNA657728 | Leaf tissue (wild vs. cultivated) | Wang et al., 2021 |
| | | GSA | CRA011652 | Seedling cotyledon, hook, hypocotyl | Li et al., 2024 |
| | Rice | SRP | SRP093806 | Aerial parts of 10-day-old seedlings | Liu et al., 2017 |
| | | PRJNA | PRJNA486213 | Mesophyll and endosperm | Dong et al., 2020 |
| | Maize | GSE / SRP | GSE120304 / SRP162341 | Inner second leaves of 6-day-old seedlings | Ricci et al., 2019 |
| | | PRJNA | PRJNA599454 | Maize ear and tassel tissues | Sun et al., 2020 |
| scATAC-seq | Soybean | GSE | GSE270392 | Leaf tissue from 10-day-old seedlings | Zhang et al., 2025 |
| | Rice | PRJNA / GSE | PRJNA1007577 / GSE252040 | Seedling/leaf tissue | Yan et al., 2024 |
| | Maize | GSE | GSE275410 | Leaf tissue from ~7-day-old seedlings | Marand et al., 2025 |

Table 3: **Data sources for Hi-C and scATAC-seq datasets.** This table provides the repository names, accession numbers, tissue information, and original publication references for all Hi-C and scATAC-seq data re-analyzed in this study. Repository acronyms: GSE = Gene Expression Omnibus, SRP = SRA Project, PRJNA = NCBI BioProject, GSA = Genome Sequence Archive.

### Supplemental Figures

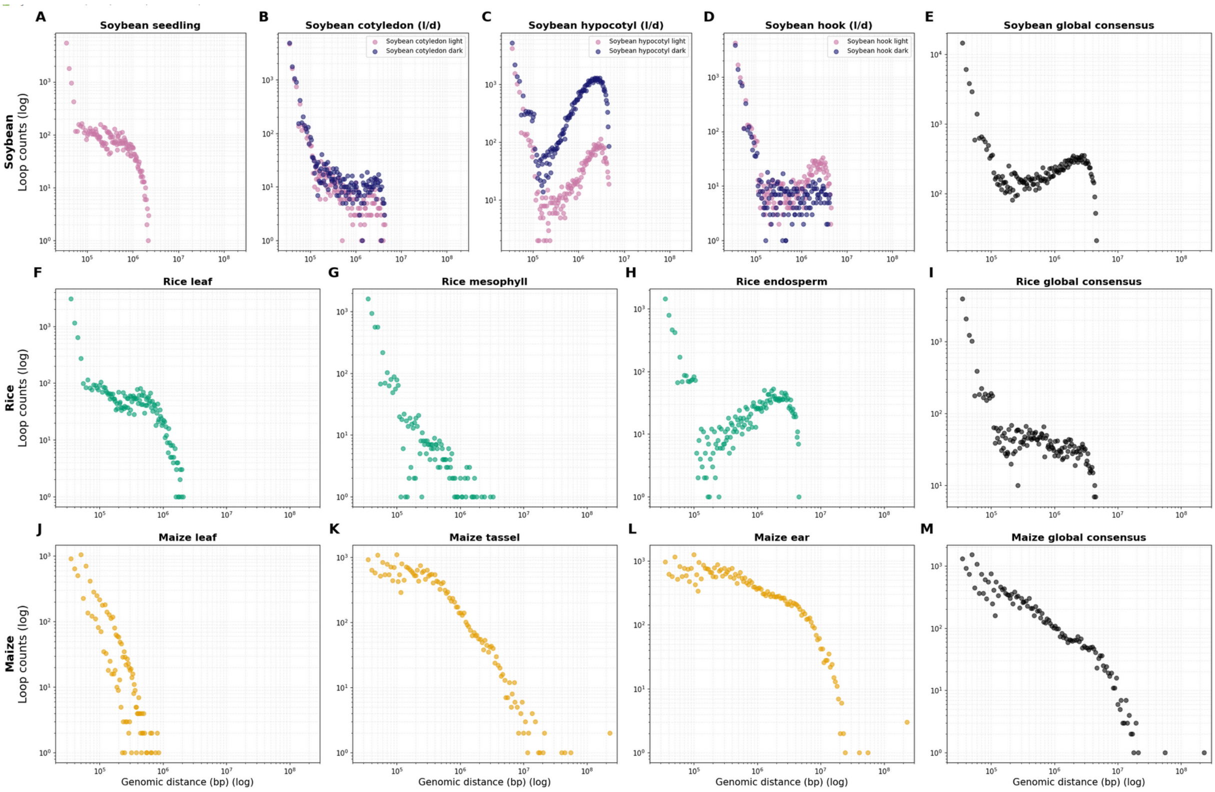


Figure 1: **Supplementary Figure S0: Raw Hi-C contact frequency scaling profiles across species and tissues.** Binned Hi-C contact counts visualized as a function of genomic distance (20 kb to 300 Mb) in log-log space. Rows represent the three species: **(A–E)** Soybean (*Glycine max*), **(F–I)** Rice (*Oryza sativa*), and **(J–M)** Maize (*Zea mays*). Individual panels correspond to specific tissues, cell types, or experimental conditions. The Global Consensus panel for each species represents a weighted integration of all available tissue datasets.

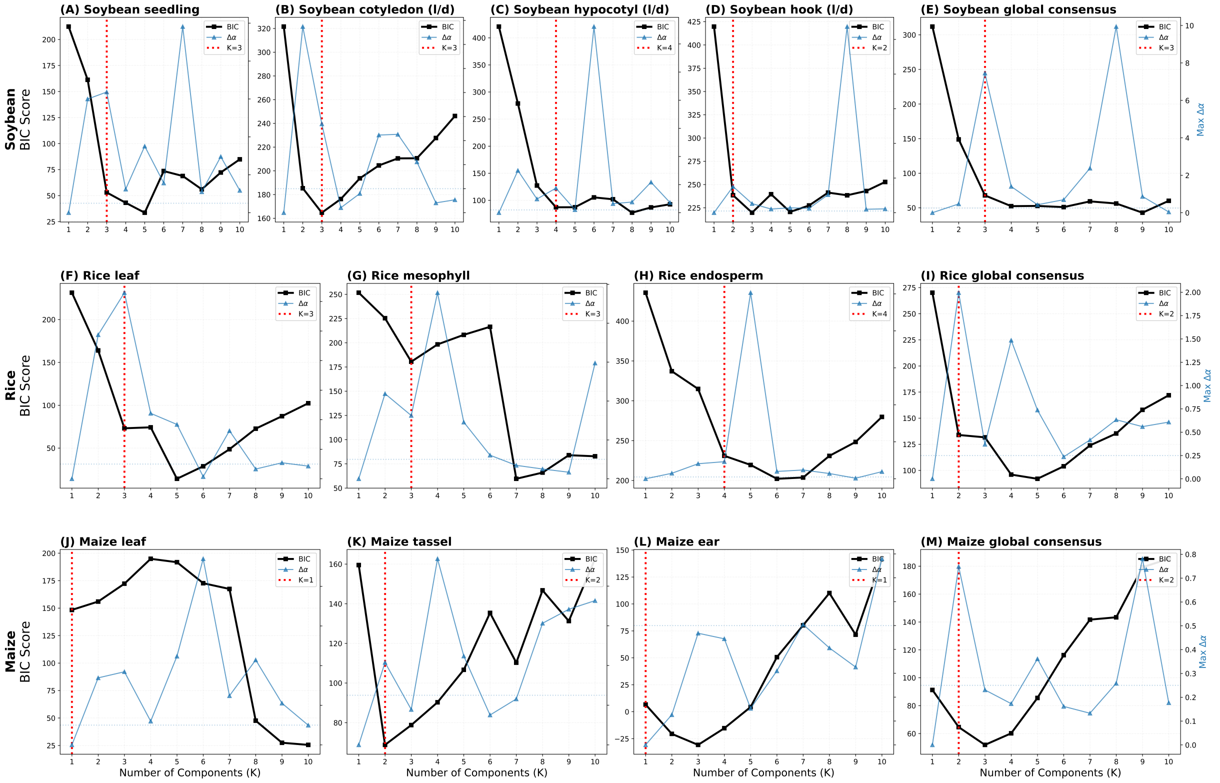


Figure 2: **Determination of the optimal number of components for the Gaussian Mixture Model.** BIC scores (black squares) and maximum change in scaling exponents ($\Delta\alpha$, blue triangles) were calculated for $K = 1$ to $K = 10$ components across all datasets in maize, rice, and soybean. Red dotted lines indicate the selected optimal $K$ for each tissue. Starting with $K = 1$, an additional compartment was only considered if BIC scores reduced significantly (requiring a $\Delta\text{BIC} > 20.0$) and the new regime simultaneously showed a significant change in the exponential decay factor $\alpha$ ($> 0.25$). In accordance with the principle of parsimony, these criteria provide the best balance between model fit and complexity to accurately capture architectural transitions while avoiding overfitting.

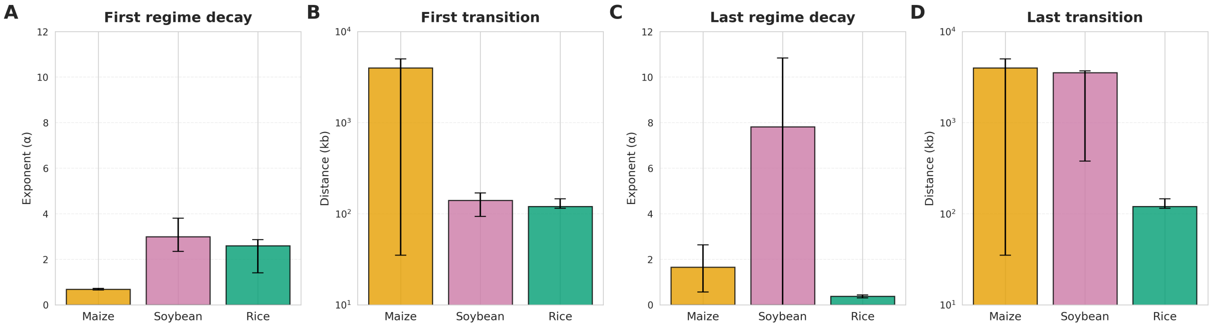


Figure 3: **Statistical robustness of GMM regimes.** Median estimates and 95% confidence intervals (CI) derived from 10,000 bootstrap iterations for **(A)** first regime decay, **(B)** first transition points, **(C)** last regime decay, and **(D)** last transition points in maize, rice, and soybean.

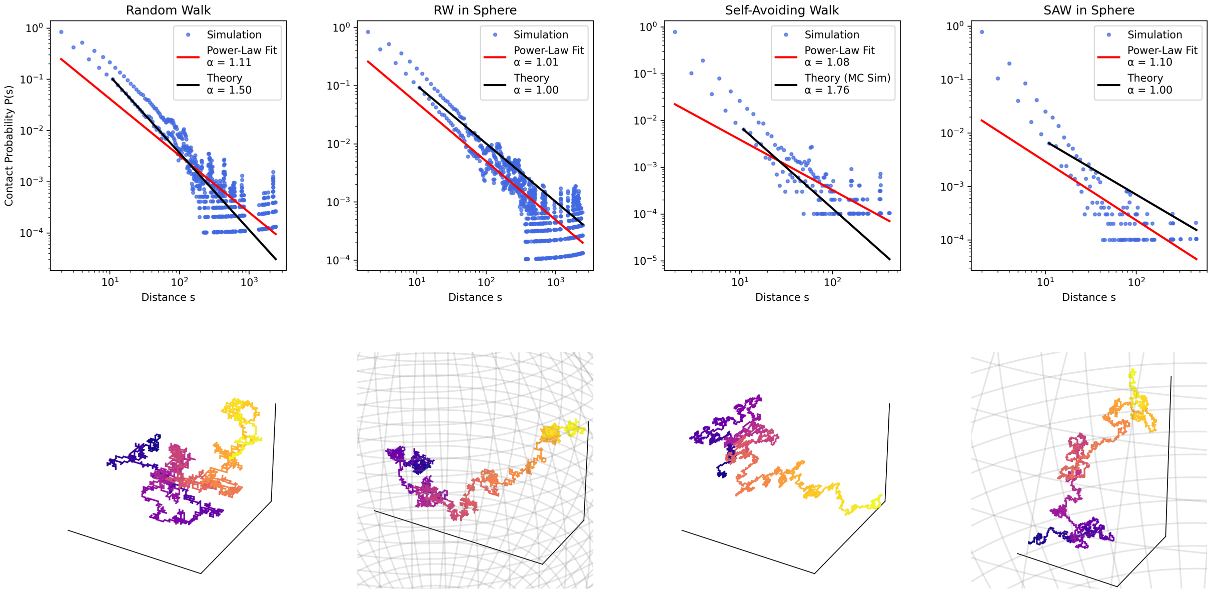


Figure 4: **Polymer simulations of chromatin scaling regimes.** Simulations ($N = 10,000$ steps) were performed using the Pivot Monte Carlo Algorithm for four models: the Ideal Chain (Random Walk, RW), the Self-Avoiding Walk (SAW), and both models under spherical confinement. The upper panels show the contact probability $P(s)$ as a function of distance $s$, which follows a power-law $P(s) \propto s^{-\alpha}$. The lower panels show a representative 3D conformation for each model. The discrepancy between these single-run $\alpha$ values and established theory is expected due to stochastic fluctuation. The key takeaway is that a lower $\alpha$ value (closer to 1.0) indicates a more compact state, as seen in the confined walk scenarios. Crucially, the failure of any of these simple, uniform models to recapitulate the multi-linear decay observed in experimental Hi-C data underscores the necessity of our multi-component approach.

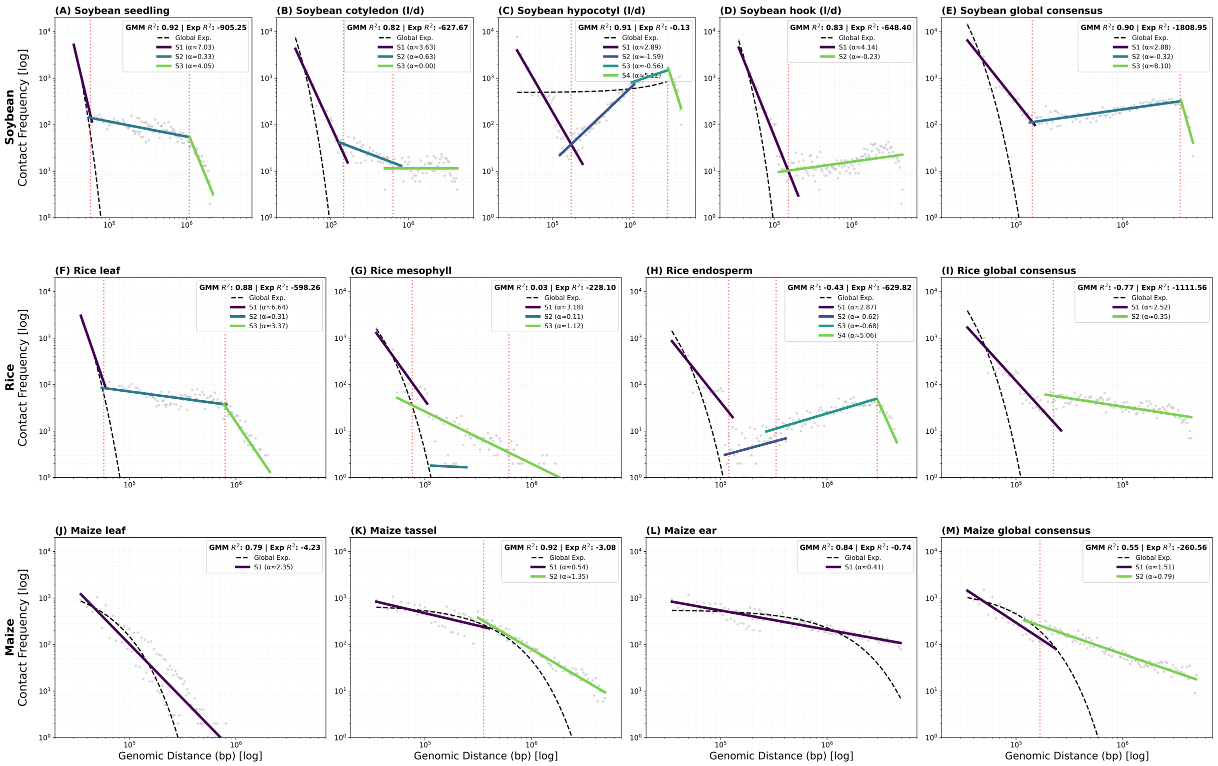


Figure 5: **Benchmarking of GMM multi-regime against global exponential fit.** Contact frequency decay profiles (gray points) for all tissues were fitted using the multi-regime GMM (colored solid lines) and a standard global exponential decay model (black dashed lines). The GMM approach performs better than exponential fit by capturing architectural transitions (red dotted lines), resulting in a minimum 11.1-fold improvement in $R^2$ accuracy.

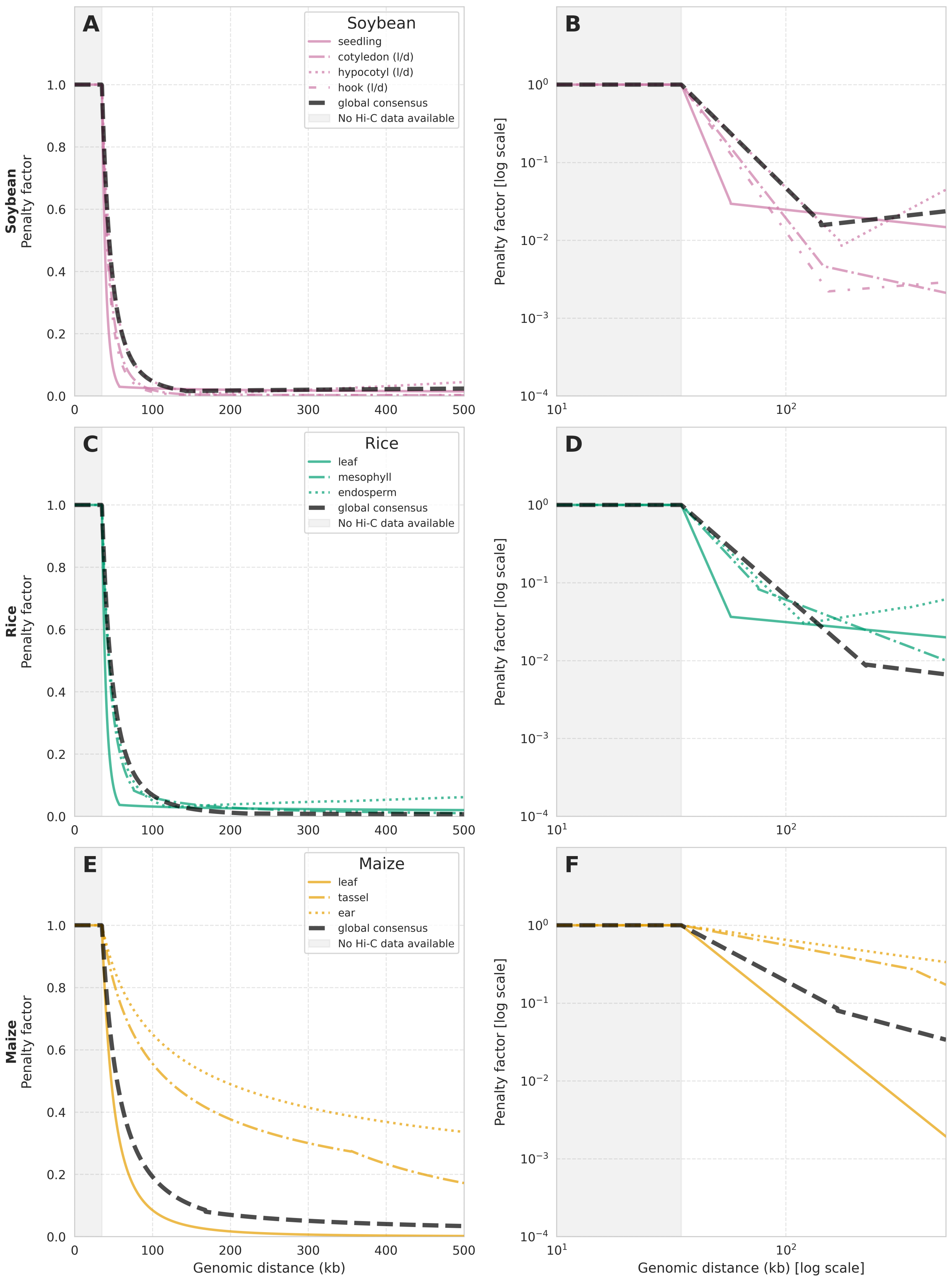


Figure 6: **Supplementary Figure S6: Tissue-specific and global-consensus penalty functions across multiple genomic scales.** Comparison of architectural decay models for **(A–B)** Soybean (*Glycine max*), **(C–D)** Rice (*Oryza sativa*), and **(E–F)** Maize (*Zea mays*). Profiles are visualized in linear (left panels) and log-log (right panels) scales to highlight short-range deviations and long-range power-law scaling, respectively. Gray shaded regions ($< 35$ kb) indicate distance ranges below the Hi-C resolution limit where data is absent. Black dashed lines represent the global-consensus penalty models, serving as a baseline against colored tissue-specific profiles, illustrating the degree of architectural conservation across different biological conditions.
